# Chromosome-scale genome assembly and annotation of the Vietnamese indica rice cultivar Khang Dan 18

**DOI:** 10.64898/2026.08.15.742683

**Authors:** Trung Q. Nguyen, Khoa H.D. Do, Thiet M. Vu, Nam V. Hoang

## Abstract

Khang Dan 18 (KD18) is an *Oryza sativa* L. subsp. *indica* rice cultivar widely cultivated in northern Vietnam and used as an experimental and breeding background in Vietnamese rice research. Although KD18 has previously been represented in low-depth population resequencing datasets, a contiguous and annotated cultivar-specific genome has not been available. Here, we report a chromosome-scale genome assembly of KD18 generated using Oxford Nanopore long-read and Illumina short-read sequencing. The 395.3-Mb assembly comprises 12 chromosome-scale pseudomolecules containing approximately 95% of the assembled sequence and 99.6% of the predicted protein-coding genes. The assembly showed 97.2% BUSCO completeness, an average Merqury quality value of 46 and a long terminal repeat assembly index of 13.21. A total of 56,546 protein-coding genes representing 71,237 transcripts were predicted, with 99% BUSCO and 98.68% OMArk completeness. These statistics are similar to those of other high-quality genome assemblies that were recently published for different Asian rice cultivars, therefore providing a cultivar-specific genomic resource for research involving KD18 and KD18-derived materials.

## BACKGROUND & SUMMARY

Asian rice (*Oryza sativa* L.) is one of the world’s most important staple crops and encompasses extensive genetic diversity across cultivated varieties and breeding germplasm^1–4^. The availability of high-quality rice genome assemblies has substantially advanced studies of genome evolution, gene function, and agronomically important traits^2,5,6^. The genome of the japonica rice cultivar Nipponbare has served as a major reference for rice genetics and genomics, while chromosome-scale and telomere-to-telomere assemblies of additional japonica and indica cultivars have revealed considerable sequence and structural variation among rice genomes^7–11^. The observations highlight the value of cultivar-specific genomic resources, particularly for experimental and breeding materials that are genetically distinct from commonly used reference cultivars. As the demand to improve rice productivity and resilience continues to increase under the pressures of population growth and climate change, comprehensive genomic resources are also becoming increasingly important for accelerating crop improvement^12,13^. Expanding the collection of high-quality genome assemblies from genetically diverse indica rice germplasm is therefore essential for comparative genomics, gene discovery, and the studies of molecular basis of agronomically important traits.

*Oryza sativa* L. subsp. *indica* cultivar Khang Dan 18 (hereafter, KD18) is an elite rice cultivar widely cultivated in northern Vietnam. KD18 is valued for its relatively short growth duration, stable yield performance, broad environmental adaptation, and suitability for intensive rice production. The cultivar typically completes its growth cycle within approximately 95 days and has been reported to produce an average grain yields of 5-5.5 t ha⁻¹^14^. Owing to these agronomic characteristics and its adaptability to experimental cultivation, KD18 has also been widely used as a genetic background in Vietnamese rice breeding and functional studies, including genetic transformation, genome editing, gene characterization, and investigations of plant responses to environmental stresses^15^. The frequent use of KD18 in experimental studies creates a practical need for an accurate cultivar-specific genome sequence to support gene identification, primer and guide-RNA design, genomic comparison, and reference-based analysis of KD18-derived materials.

Although, KD18 has previously been included in population-scale resequencing studies of Vietnamese rice germplasm^16^, these data were generated at relatively low sequencing depth and primarily supported variant discovery and population-genetic analyses. Such resequencing data do not provide a contiguous and structurally annotated cultivar-specific genome. In addition, sequence information generated from KD18-based functional studies has largely remained restricted to individual genes or genomic loci. Consequently, a high-quality assembled and annotated KD18 genome suitable for cultivar-specific genomic analyses and experimental design has remained unavailable.

Here, we generated approximately 25 Gb of Oxford Nanopore long-read data (62× genome coverage) and 60 Gb of Illumina paired-end (PE) short-read data (150×) from KD18 and constructed a 395.3-Mb genome assembly. The final *de novo* assembly was organized into 12 chromosome-scale pseudomolecules by reference-guided scaffolding against high-quality indica rice genomes. We provide the assembled genome, protein-coding gene annotation models, coding and protein sequences, repeat annotations, and associated raw sequencing data. The assembly quality was evaluated using k-mer-based consensus assessment, conserved gene completeness, repeat-space contiguity, independent previous published KD18 resequencing data, and the syntenic concordance with other high-quality genomes. This dataset provides a cultivar-specific genomic resource for studies involving KD18 and KD18-derived experimental and breeding materials.

## METHODS

### Plant materials, sample preparation and genomic DNA extraction

Plant materials used in this study were derived from the rice cultivar KD18 (**Fig. 1A**) which was maintained and phenotypically characterized over multiple growing seasons at the Vietnam National University of Agriculture (VNUA), Hanoi, Vietnam. For whole genome sequencing (WGS), KD18 seeds from the maintained germplasm line described above were germinated and seedling were grown under lab conditions for two weeks. Young leaves were collected, and high-molecular weight (HMW) genomic DNA was isolated using a modified CTAB method as described by Kang et al., 2023^17^. DNA integrity was checked on agarose 1% electrophoresis. DNA purity and concentration were measured by using a NanoDrop One Spectrophotometer (Thermo Fisher Scientific, Waltham, MA, USA) and Qubit fluorometer (Thermo Fisher Scientific), respectively.

**Figure 1.**
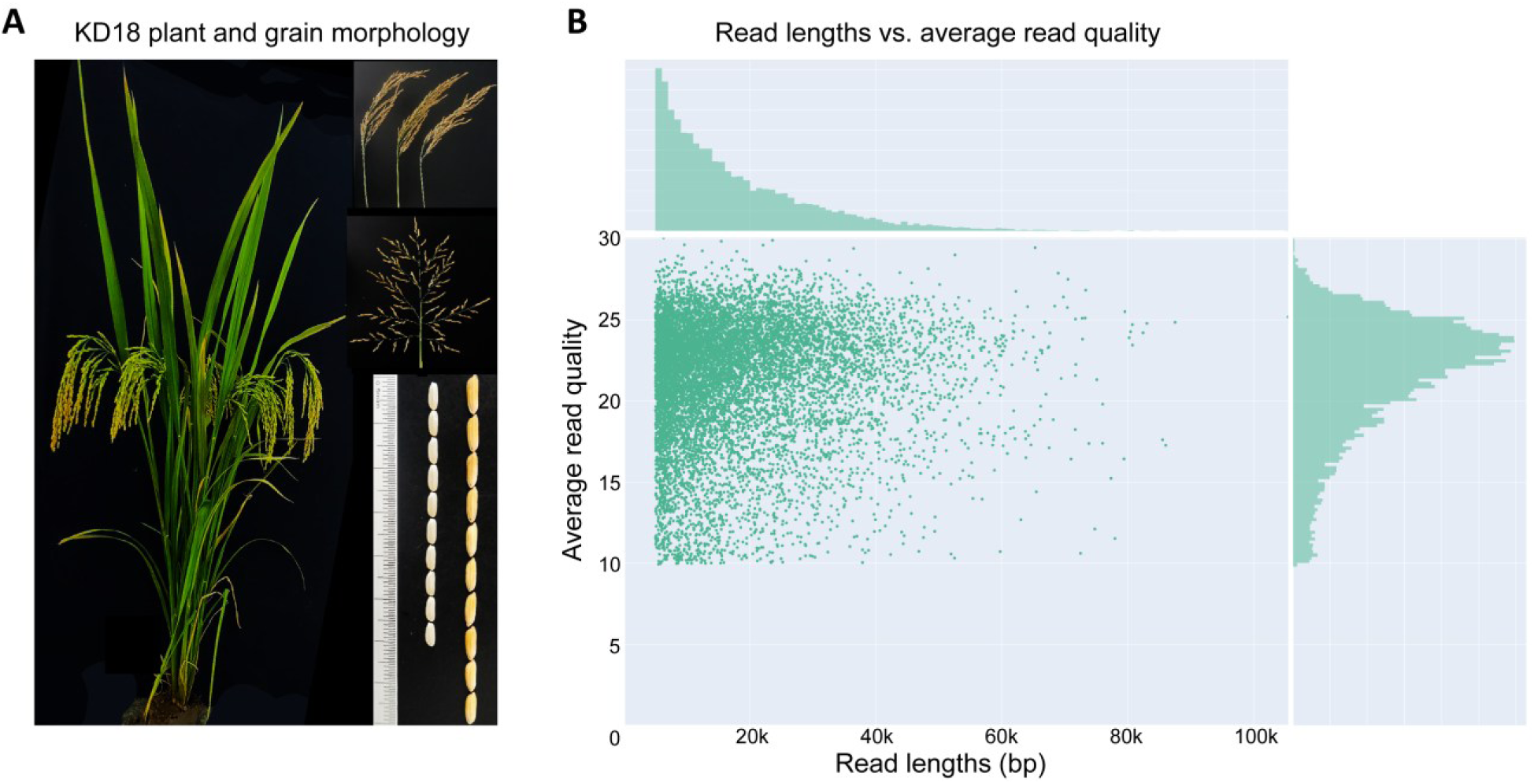
Sample preparation and data generation of rice cultivar KD18 (*O. sativa* L. subsp. indica). **(A)** Plant and grain morphology, including whole-plant architecture, panicles, flowers and grains. **(B)** A summary of read length vs. read quality of a total of 25 Gb of trimmed ONT sequencing data (1,395,516 reads, N50 of 23 kb) generated in this study.

### Oxford Nanopore long-read sequencing

An aliquot of 3 ug of HMW genomic DNA was used for library preparation following the SQK-LSK114 kit of the Oxford Nanopore Technologies (ONT, Oxford, UK). We generated a library with size selection of a >25 kb threshold using the short fragment eliminator kit (EXP-SFE001, ONT). The library was sequenced on a PromethION P2 solo platform (ONT, Oxford, UK) using an R10.4.1 flow cell, and base-calling was performed using the built-in dorado software with default parameters and “*--min-qscore 10*”. Adapters were checked and removed using PoreChop v0.2.4 (https://github.com/rrwick/porechop). The adapter-free raw ONT data was then trimmed for quality (min Q10) and length (min 5 kb) using Chopper v0.8.0 (https://github.com/wdecoster/chopper) before downstream analyses. Read length and quality distribution was assessed using Nanoplot and NanoComp from the NanoPack v1.1^18^. After filtering, the final trimmed ONT data consisted of 25 Gb, with 1,395,516 reads and a read N50 of 23 kb (**Fig. 1B**), representing roughly 62× genome coverage of the estimated KD18 genome size.

### Illumina short-read sequencing

Approximately 0.4 µg of genomic DNA was used for Illumina library preparation and PE sequencing (2×150 bp) on a NovaSeq 6000 platform at NOVOGENE (Singapore). Adapter sequences, low-quality and duplicate reads were trimmed using Fastp v1.3.5^19^ with options “*--detect_adapter_for_pe -- trim_poly_x -q 20 -l 75*”, and Trimmomatic v0.41^20^ with options: “*ILLUMINACLIP:TruSeq3-PE.fa:2:30:7:1:true LEADING:3 TRAILING:3 SLIDINGWINDOW:4:30; MINLEN:75.*” Read quality before and after trimming was assessed by FastQC v0.12.1 (https://github.com/s-andrews/fastqc). Approximately 405.8 million clean PE reads (60 Gb, Phred Q score ≥30) were obtained which represented around 150× genome coverage of the estimated KD18 genome size.

### Genome size estimation of KD18

To estimate the expected genome size of cultivar KD18, we first used the total 405.8 million Illumina PE WGS data and GenomeScope v2.0^21^. The k-mer distribution was generated by KMC v3.2.4^22^ with a k-mer ranging from 21 to 141, and the “*-cx1000000*” option to account for the high-frequency k-mers derived from repetitive elements in the genome. The genome size estimation for KD18 cultivar was around 399 Mb (**Fig. 2**), which is in line with other reported results for indica rice genomes^7–11^. Additionally, we also utilized a total of 25 Gb long-read ONT data to predict the genome size of KD18 by ONT-read-check v0.2.0 (https://github.com/SamD28/ONT-read-check.git). The ONT-based prediction was 440 Mb, which is a bit larger than expected genome size. Nevertheless, the results indicate that KD18 possesses a typical genome size of an indica rice.

**Figure 2.**
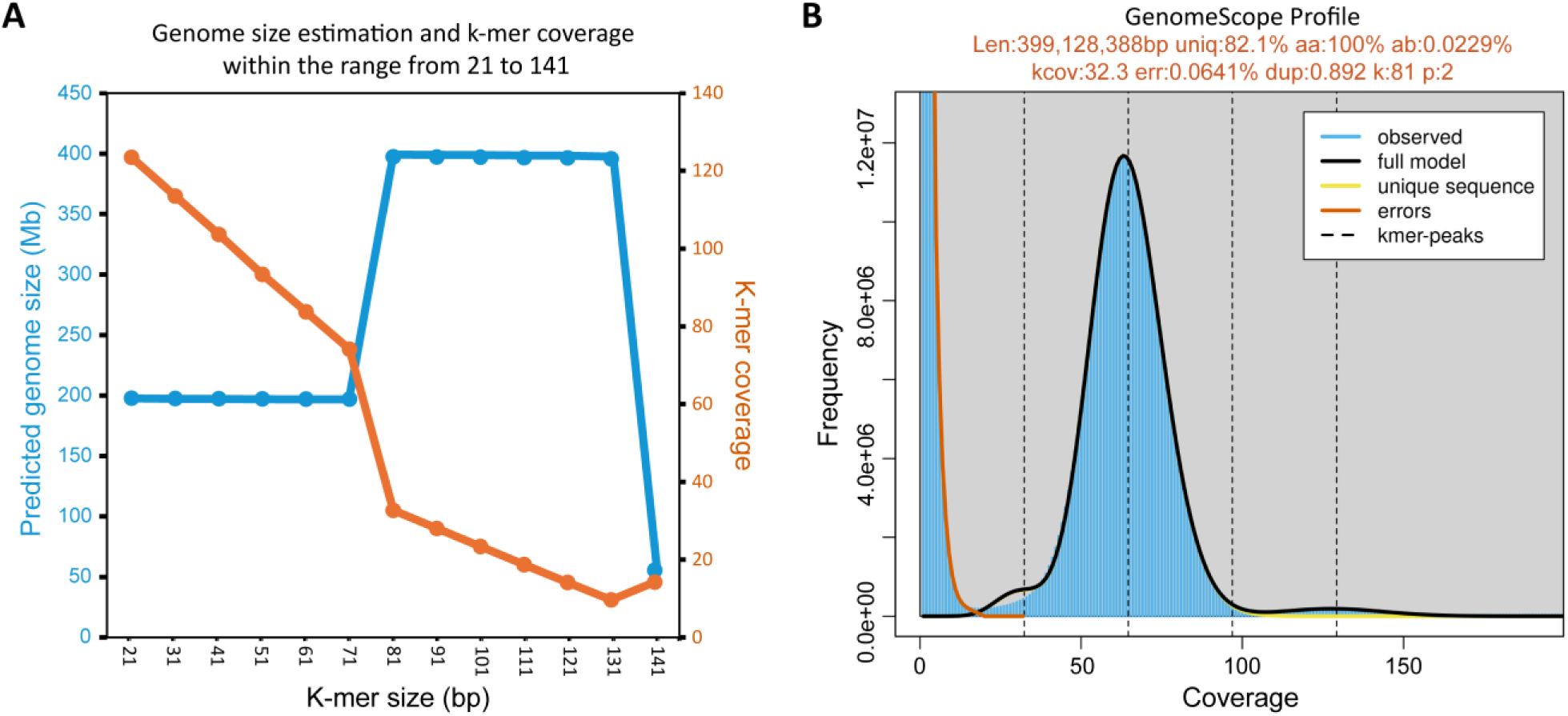
Genome size prediction by GenomeScope 2.0 for rice cultivar KD18 (*O. sativa* L. subsp. indica) based on Illumina data. **(A)** Genome size prediction using a range of k-mer from 21 to 141. **(B)** GenomeScope profile of KD18 genome predicted using k-mer of 81 (k). The predicted genome size of 399 Mb is shown in the figure, together with k-mer coverage (kcov), heterozygosity (ab%) and ploidy level (p).

### *De novo* genome assembly and chromosome-scale scaffolding

The final trimmed ONT reads of a minimum length of 5 kb and a minimum quality score of Q10 were used for genome assembly by Flye^23^ with the options “*--nano-raw --genome-size 399m --scaffold -- iterations 2 --no-alt-contigs*”. After two rounds of long-read polishing within the Flye pipeline, we obtained a draft assembly with a size of 395.3 Mb (accounting for 99% of predicted genome size) consisting of 694 sequences. The draft genome was further polished using 405.8 million Illumina PE short reads by NextPolish v1.4.1^24^ to correct base-level substitution and small INDELs errors. The polished assembly showed a completeness of 97.2% by the Benchmarking Universal Single-Copy Orthologs (BUSCO) v6.1.0^25^ running with the options “-*m genome --metaeuk -l poales_odb12*” in which MetaEuk^26^ was employed to search genes against 6,282 Poales conserved orthoglogs. This completeness score is comparable with the other recently published chromosome-level genome assemblies for rice^7–11^ (**Table 1**).

**Table 1.** A summary statistics and comparison of KD18 genome assembly.

|  | **KD18 draft** | **KD18**<br>**based on**<br>**MH63\*** | **KD18**<br>**based on**<br>**ZS97\*** | **KD18**<br>**based on**<br>**T197\*** | **KD18**<br>**based on 3**<br>**genomes\*** | **Osa1_v7**<br>**(Osa**<br>**japonica)** | **AGIS1**<br>**(Osa**<br>**japonica)** | **T197 (Osa**<br>**indica)** | **MH63**<br>**(Osa**<br>**indica)** | **ZS97 (Osa**<br>**indica)** |
| --- | --- | --- | --- | --- | --- | --- | --- | --- | --- | --- |
| **Number of scaffolds** | 694 | 149 | 130 | 152 | 115 | 14 | 12 | 12 | 12 | 12 |
| Number of contigs | 707 | 707 | 707 | 707 | 707 | 298 | 12 | 12 | 12 | 12 |
| **Total length** | **395,256,163** | **395,310,663** | **395,312,563** | **395,310,363** | **395,314,063** | **374,471,240** | **385,711,369** | **395,084,276** | **395,765,488** | **391,561,630** |
| Percent gaps | 0.00% | 0.0140% | 0.0150% | 0.0140% | 0.0150% | 0.0440% | 0.00% | 0.00% | 0.00% | 0.00% |
| Scaffold N50 (Mb) | 1 | 29 | 29 | 29 | 29 | 29 | 31 | 31 | 31 | 32 |
| **Complete BUSCOs (C)**<br>**in %** | **97.2** | **97.2** | **97.2** | **97.2** | **97.2** | **97.0** | **97.2** | **97.1** | **97.1** | **97.2** |
| **Complete BUSCOs (C)** | **6,105** | **6,104** | **6,104** | **6,104** | **6,104** | **6,093** | **6,108** | **6,102** | **6,098** | **6,104** |
| Complete and single-copy<br>BUSCOs (S) | 6,030 | 6,033 | 6,032 | 6,032 | 6,033 | 6,023 | 6,043 | 6,035 | 6,029 | 6,040 |
| Complete and duplicated<br>BUSCOs (D) | 75 | 71 | 72 | 72 | 71 | 70 | 65 | 67 | 69 | 64 |
| Fragmented BUSCOs (F) | 44 | 46 | 46 | 46 | 46 | 52 | 46 | 52 | 51 | 49 |
| Missing BUSCOs (M) | 133 | 132 | 132 | 132 | 132 | 137 | 128 | 128 | 133 | 129 |
| Total BUSCO groups<br>searched | 6,282 | 6,282 | 6,282 | 6,282 | 6,282 | 6,282 | 6,282 | 6,282 | 6,282 | 6,282 |

\*Different versions scaffolded based on different reference genomes using RagTag <sup>27</sup>.
*Osa* denotes *Oryza sativa*.
Osa1\_v7 and AGIS1 are japonica rice cultivar Nipponbare.
BUSCO based on 6,282 Poales conserved orthoglogs (odb12).

To scaffold the draft genome assembly, we employed RagTag v2.1.0^27^ using the *“scaffold”* function based on three chromosome-level assemblies for the indica rice, including Minghui 63 (MH63)^8^, Zhenshan 97 (ZS97)^8^ and Kato T197 (T197)^7^. The results showed that chromosome-level assemblies could be obtained for KD18 using this approach, in which, for all cases, we were able to obtain 12 main scaffolds that represent around 95% of the original draft assembly (**Fig. 3A** and **Table 1**). Since our results showed that these resultant assemblies were very similar, we selected the assembly scaffolded using the MH63 genome for further analyses because it contained a lower percentage of gaps.

**Figure 3.**
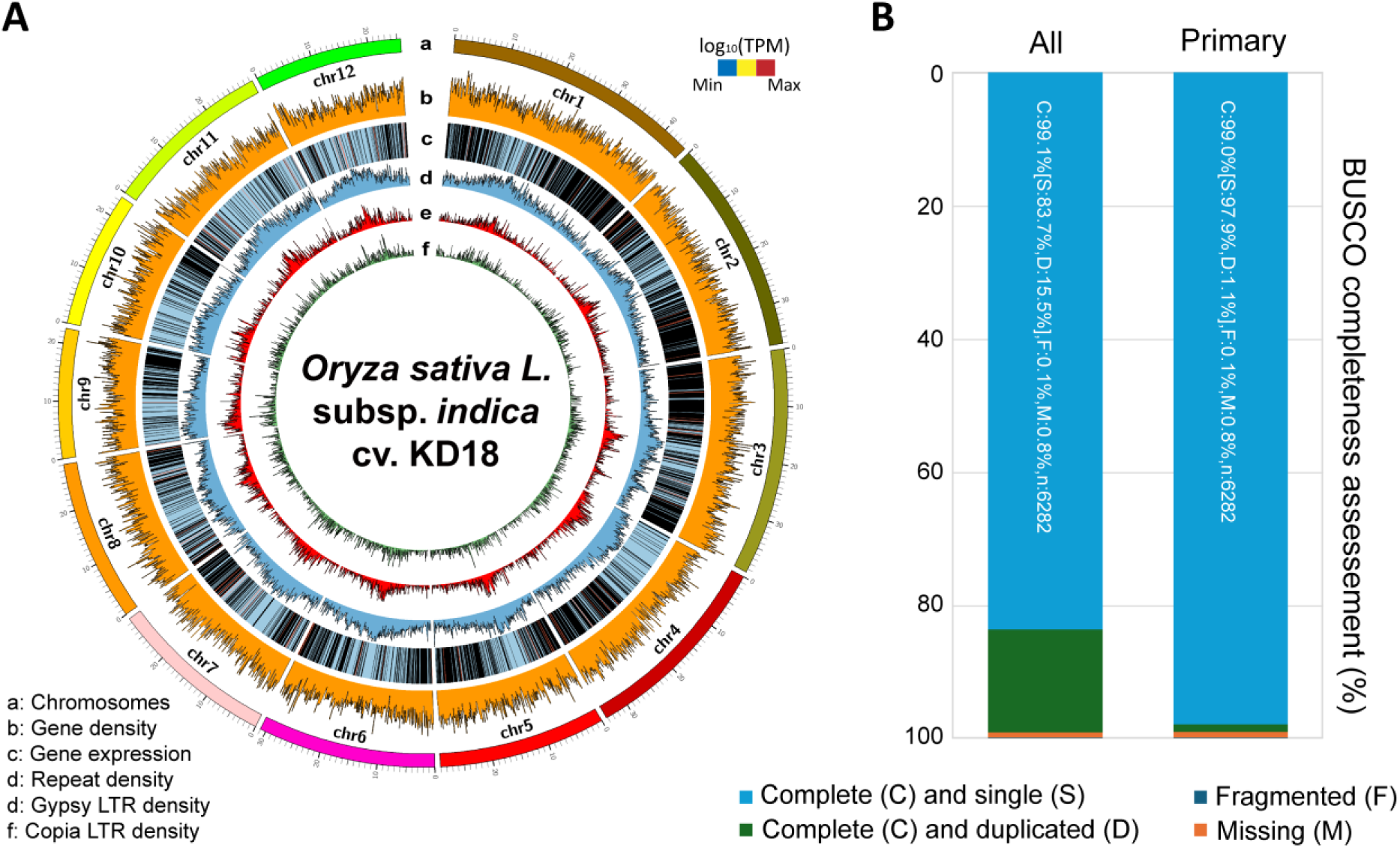
The assembly and annotation of the genome of rice cultivar KD18 (*O. sativa* L. subsp. indica). **(A)** A Circos plot showing different features mapped onto the 12 pseudo-chromosomes of the KD18 genome, including chromosome length (track **a**), gene density (track **b**), gene expression (track **c**), repeat distribution (track **d**), LTR *Gypsy* repeat class distribution (track **e**) and LTR *Copia* repeat class distribution (track **f**). Length in Mb. Distributions were estimated for each window of 100 kb. For gene expression, RNA-seq reads were mapped onto the KD18 genome by HISAT2 v2.2.1^29^ with default settings and gene expression levels (TPM) were estimated using StringTie v3.0.3^30^. Gene expression was quoted as log10[transcripts TPM]. The plot was created using Circos^31^. (**B**) BUSCO completeness score of the annotation of the KD18 genome, including all transcripts and primary transcripts.

The final genome assembly for KD18 has 12 pseudo-chromosomes and 137 small unplaced scaffolds, with a length of 395.3 Mb and an N50 of 29 Mb (**Fig. 3A** and **Table 1**). This chromosome-level assembly achieved a long terminal repeat assembly index (LAI) of 13.21 through LTR_retriever v3.0.5^28^, while retained the same total BUSCO completeness of 97.2% after scaffolding. The LAI value also indicates that the genome assembly is of reference genome-level quality (10 ≤ LAI < 20)^28^. Collectively, our *de novo* genome assembly of KD18 is comparable in quality with recently published high-quality rice genomes^7–11^.

### Genome repeat and gene annotation of KD18

Repetitive and transposable elements in the KD18 genome were masked with RepeatModeler v2.0.9/RepeatMasker v4.1.2^32^. Firstly, the *ab initio* prediction program RepeatModeler was employed to build a *de novo* repeat library based on the KD18 genome. Then, using a custom library that consisted of *de novo* identified repeats, Dfam v4.0 and RepBase RepeatMasker Edition 20181026 as database, RepeatMasker was run to discover and classify repetitive elements in the genome. A total of 208 Mb (52.71%) were classified as repetitive elements in the KD18 genome, with 24.57% retroelements (i.e., SINEs, LINEs and long terminal repeat/LTR) and 12.54% DNA transposons (**Fig. 3A** and **Table 2**). Among the LTR elements, two main classes, Gypsy/DIRS1 and Ty1/Copia, accounted for 18.39% and 3.52%, respectively.

**Table 2.** Repeat masking result of the KD18 genome.

| Repeat class |  |  | Number of elements | Length occupied (bp) | Percentage of sequence (%) |
| --- | --- | --- | --- | --- | --- |
| Retroelements |  |  | 90,631 | 97,108,263 | 24.57 |
|  | SINEs: |  | 5,241 | 767,318 | 0.19 |
|  | Penelope |  | 37 | 20,298 | 0.01 |
|  | LINEs: |  | 13,360 | 5,860,151 | 1.48 |
|  |  | CRE/SLACS | 0 | 0 | 0 |
|  |  | L2/CR1/Rex | 0 | 0 | 0 |
|  |  | R1/LOA/Jockey | 0 | 0 | 0 |
|  |  | R2/R4/NeSL | 2 | 77 | 0 |
|  |  | RTE/Bov-B | 88 | 9,561 | 0 |
|  |  | L1/CIN4 | 12,439 | 5,573,130 | 1.41 |
|  | LTR elements: |  | 72,030 | 90,480,794 | 22.89 |
|  |  | BEL/Pao | 52 | 6,669 | 0 |
|  |  | Ty1/Copia | 17,874 | 13,902,498 | 3.52 |
|  |  | Gypsy/DIRS1 | 42,471 | 72,713,954 | 18.39 |
|  |  | Retroviral | 392 | 103,332 | 0.03 |
| DNA transposons |  |  | 169,314 | 49,558,151 | 12.54 |
|  | hobo-Activator |  | 22,498 | 5,007,181 | 1.27 |
|  | Tc1-IS630-Pogo |  | 22,217 | 3,478,185 | 0.88 |
|  | En-Spm |  | 0 | 0 | 0 |
|  | MuDR-IS905 |  | 0 | 0 | 0 |
|  | PiggyBac |  | 0 | 0 | 0 |
|  | Tourist/Harbinger |  | 28,901 | 6,143,056 | 1.55 |
|  | Other (Mirage, P-element, Transib) |  | 22 | 2,699 | 0 |
| Rolling-circles |  |  | 46,231 | 11,774,427 | 2.98 |
| Unclassified: |  |  | 185,854 | 45,242,545 | 11.44 |
| Total interspersed repeats: |  |  |  | 191,908,959 | 48.55 |
| Small RNA: |  |  | 4,681 | 820,714 | 0.21 |
| Satellites: |  |  | 464 | 65,632 | 0.02 |
| Simple repeats: |  |  | 84,586 | 3,916,354 | 0.99 |
| Low complexity: |  |  | 8,531 | 414,663 | 0.1 |
| Total genome |  |  |  | 395,310,663 | 100 |
| Total bases masked |  |  |  | 208,368,851 | 52.71 |

For *de novo* gene prediction, the BRAKER pipeline v3.0.8^33^ was employed to generate a hint file based on RNA-seq data and protein sequences publicly available for Asian rice. For RNA-seq hints, we obtained previously published RNA-seq data for indica rice from several studies^7,34–37^, which include samples representing key organs and tissues such as leaf, root, stem, flower, fruit peel and flesh of the rice plants. Additionally, the Viridiplantae protein dataset derived from the OrthoDB v11^38^ was combined with the rice proteins obtained from recently published high-quality indica and japonica rice genomes^7–11^ to generate protein hints. After that, GeneMark-ETP/EP+^39^ used the hints to predict initial high-confidence gene models and training genes that were subsequently used by AUGUSTUS v4.0.0^40^ to train and predict genes across the whole genome. We then used PASA v2.5.3^41^ to update gene models that were not in agreement with the RNA-seq data and to add untranslated regions (UTR) to the BRAKER-derived annotation. Finally, a total of 56,546 gene models (71,237 transcripts) were predicted for the KD18 genome that displayed a BUSCO score of 99% based on 6,282 Poales conserved orthoglogs, and 98.68% proteome completeness score by OMArk v0.5.0^42^ based on 13,374 conserved orthologs of the Poaceae family (**Fig. 3B**). Among the total predicted genes, 99.6% (56,315) were anchored onto the 12 pseudo-chromosomes of the KD18 assembly.

### Gene functional annotation

A total of 98.31% of the 56,546 KD18 predicted proteins matched at least one of the public protein databases (**Table 3**), including 91.88% matching the TrEMBL/Swiss-Prot release 2026_02^43^ using Diamond BLASTP v2.0.14^44^ with the following settings “*-e 1e-5 -k 1*”, and 82.05% matching the OMA database according to OMArk^45^. Additionally, to predict protein function, we used InterProScan-5.78-109.0^46^ to blast KD18 proteins against a total of 18 databases provided with InterProScan with the options “*-iprlookup -goterms --pathways*”, and eggNOG v5^47^ with the option *“--tax_scope Viridiplantae”*. Here, we found that 89.99% of KD18 predicted proteins had at least one InterProScan annotation, 62.57% were assigned an InterPro accession, 63.3% contained a PFAM domain, 43.79% were associated with Gene Ontology (GO) terms^48^, and 25.08% were assigned to KEGG pathways^49^. Together with 99% BUSCO completeness and 98.68% OMArk completeness, these results support the completeness and broad biological consistency of the KD18 protein-coding gene annotation.

**Table 3.**
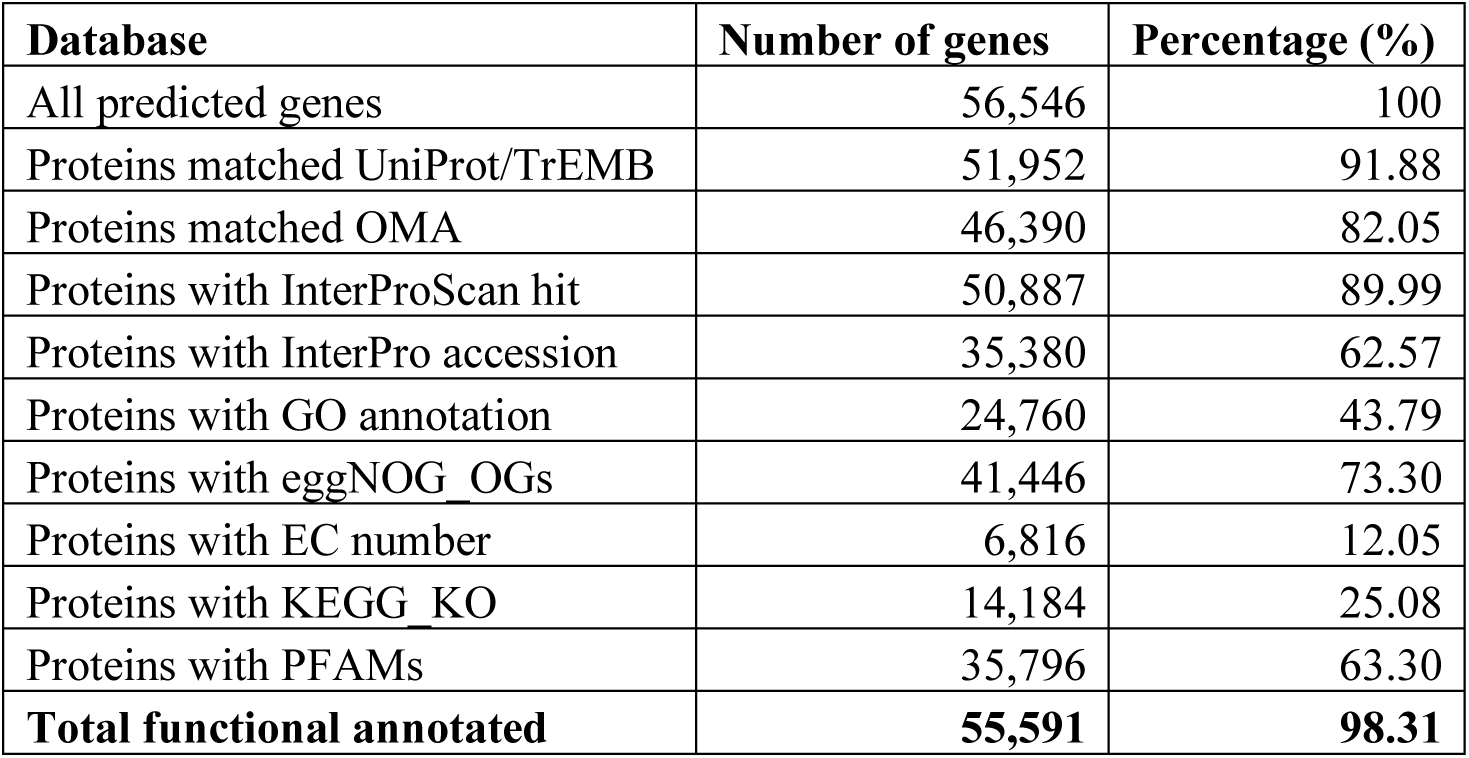
Functional annotation of the KD18 genome assembly.

### Comparative genomics based on orthogroup inference

Finally, orthogroup classification and phylogenetic analysis were conducted to evaluate the quality and evolutionary consistency of the KD18 annotated gene set. First, to infer the orthology of KD18 and other *O. sativa* genomes (i.e., indica rice IR64^37^, T197, MH63; and japonica rice AGIS1 and Zhonghua11/ZH11), primary protein sequences were used for orthogroup clustering by OrthoFinder v2.5.5^50^ with default settings. Our orthogroup clustering analysis (**Fig. 4A** and **Table 4**) found that about 95.8% of KD18 genes (54,162) were clustered in orthogroups, comparable to that of other cultivars which ranged from 93.6% (MH63) to 98.8% (IR64). The percentage of genes in species-specific orthogroups of the KD18 genome was 2.6 (1,469 genes), while the percentage of unassigned genes was 4.2% (2,384 genes). In general, these values, together with the number of orthogroups per genome, were in the range observed for other genomes included in the analysis (**Table 4**).

**Figure 4.**
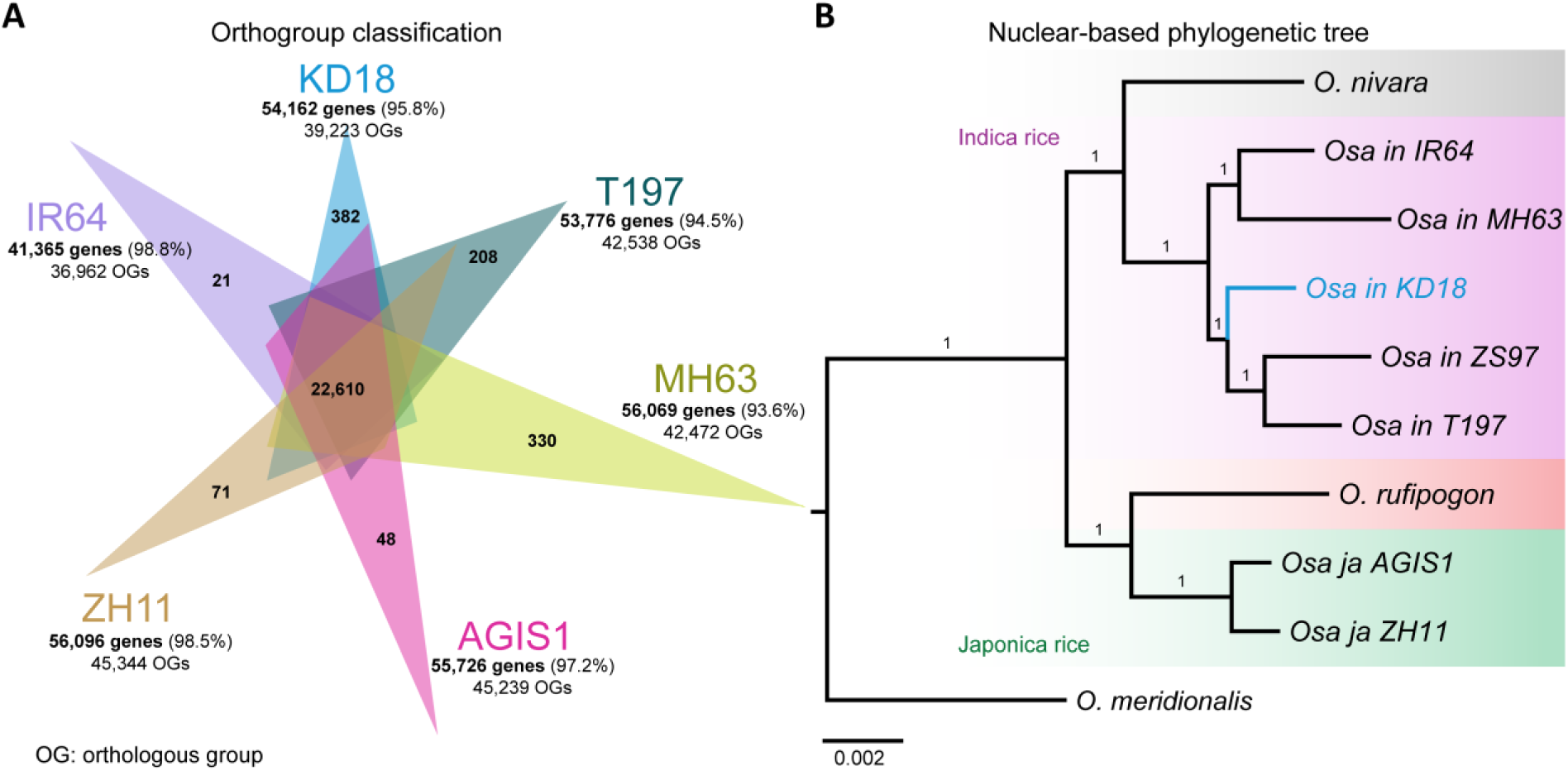
Orthogroup classification and phylogenetic analysis of the KD18 genome. (**A**) Venn diagram showing shared and unique orthogroups among different selected rice genomes. These include indica rice KD18 (this study), IR64^37^, MH63^8^, T197^7^, and japonica rice Nipponbare (AGIS1)^9^ and ZH11^11^. The numbers of genes in orthogroups and orthogroups are shown for each genome (details are in **Table 4**). Percentages were calculated based on the total genes annotated in each selected genome. OG denotes orthogroup. Venn diagram was created by Evenn tool (http://www.ehbio.com/test/venn). **(B)** Nuclear-based phylogenetic tree using 8,224 single copy genes identified by OrthoFinder amongst 10 selected rice genomes. The ZS97 was obtained from Song, et al. ^8^; while the genomes of *O. nivara*, *O. rufipogon* and *O. meridionalis* were download from ENSEMBL plant release 57 (https://ftp.ensemblgenomes.ebi.ac.uk/pub/plants/release-57/). Tree was rooted using the *O. meridionalis* genome as outgroup. Supporting values at each node are bootstrap scores (maximum is 1 or 100%). Branch length is substitutions per site.

**Table 4.** Orthologous grouping of KD18 genome with other rice genomes.

|  | **KD18**<br>**(*Osa indica*)** | **IR64**<br>**(*Osa indica*)** | **T197 (*Osa***<br>***indica*)** | **MH63**<br>**(*Osa indica*)** | **AGIS1**<br>**(*Osa***<br>***japonica*)** | **ZH11**<br>**(*Osa***<br>***japonica*)** |
| --- | --- | --- | --- | --- | --- | --- |
| Number of genes | 56,546 | 41,880 | 56,908 | 59,903 | 57,359 | 56,948 |
| Number of genes in orthogroups | 54,162 | 41,365 | 53,776 | 56,069 | 55,726 | 56,096 |
| Number of unassigned genes | 2,384 | 515 | 3,132 | 3,834 | 1,633 | 852 |
| Percentage of genes in orthogroups (%) | 95.8 | 98.8 | 94.5 | 93.6 | 97.2 | 98.5 |
| Percentage of unassigned genes (%) | 4.2 | 1.2 | 5.5 | 6.4 | 2.8 | 1.5 |
| Number of orthogroups containing genome | 39,223 | 36,962 | 42,538 | 42,472 | 45,239 | 45,344 |
| Percentage of orthogroups containing genome (%) | 65.7 | 61.9 | 71.2 | 71.1 | 75.8 | 75.9 |
| Number of genome-specific orthogroups | 382 | 21 | 208 | 330 | 48 | 71 |
| Number of genes in genome-specific orthogroups | 1,469 | 76 | 764 | 1,057 | 131 | 279 |
| Percentage of genes in genome-specific orthogroups (%) | 2.6 | 0.2 | 1.3 | 1.8 | 0.2 | 0.5 |

*Osa* denotes *O. sativa*.
AGIS1 is japonica rice cultivar Nipponbare.

Next, to ascertain the phylogenetic placement of the KD18 cultivar, in relation to other Asian rice, we also reconstructed a maximum likelihood (ML) phylogenetic tree of single-copy genes (SCGs) identified across rice cultivars of which the high-quality genomes are available. Among 10 rice cultivars included in our analysis, OrthoFinder identified 8,224 nuclear SCGs which were subsequently used for tree reconstruction by MAFFT^51^ and IQ-TREE^52^ within the pipeline with the options “-*M msa -A mafft -T iqtree*” to infer ML gene trees from multiple sequence alignment. Our nuclear-based species tree suggested that KD18 was grouped closer to two Chinese indica cultivars T197 and ZS97, and separated from MH63 and IR64 cultivars (**Fig. 4B**). Taken together, the orthogroup classification and phylogenetic analyses indicate that the predicted gene set is evolutionarily consistent with those of other rice genomes, providing additional confidence in the quality of the gene annotation.

## DATA RECORDS

The raw sequencing data generated for this study have been deposited in NCBI SRA database under BioProject PRJNA1491377, including BioSamples SAMN61953581 for ONT data and SAMN61912143 for Illumina data. The final genome assembly and annotation (v1.0) of cultivar KD18 described in this paper can be downloaded from Figshare at https://doi.org/10.6084/m9.figshare.33088043.

The dataset includes the following files:

- Osa_in_KD18_genome.fasta: genome assembly of cultivar KD18.
- Osa_in_KD18_genome.gff3: genome annotation of cultivar KD18.
- Osa_in_KD18_genome_repeats.gff3: repeat elements annotation of cultivar KD18 genome.
- Osa_in_KD18.cds_all.fasta: all extracted CDS sequences of of cultivar KD18 genome.
- Osa_in_KD18.protein_all.fasta: all extracted protein sequences of of cultivar KD18 genome.
- Osa_in_KD18.cds_longest.fasta: longest extracted CDS sequences of of cultivar KD18 genome.
- Osa_in_KD18.protein_longest.fasta: longest extracted CDS sequences of of cultivar KD18 genome.

## TECHNICAL VALIDATION

We performed three additional analyses to validate the KD18 genome assembly, including a k-mer-based consensus quality assessment, a read mapping using cultivar-specific and cross-cultivar resequencing data, and a syntenic analysis with other high-quality genomes.

### Genome assembly quality assessment using Merqury

Merqury v1.4.1^53^ was used to evaluate the consensus base-level accuracy and completeness of the KD18 genome assembly using a reference-free, k-mer-based approach. This approach provides an independent assessment of assembly accuracy by comparing the k-mer content of the assembly with that of the high-quality sequencing reads. First, a k-mer database was generated from 405.8 million clean Illumina PE reads of the KD18 cultivar using Meryl v1.4.1, and then compared with the genome assembly to estimate consensus quality value (QV) and k-mer completeness. The final KD18 genome assembly achieved a Merqury average QV of 46 (ranging from 42 to 49.2) and a k-mer completeness of 99.3% (**Table 5**), indicating high consensus accuracy and completeness.

**Table 5.**
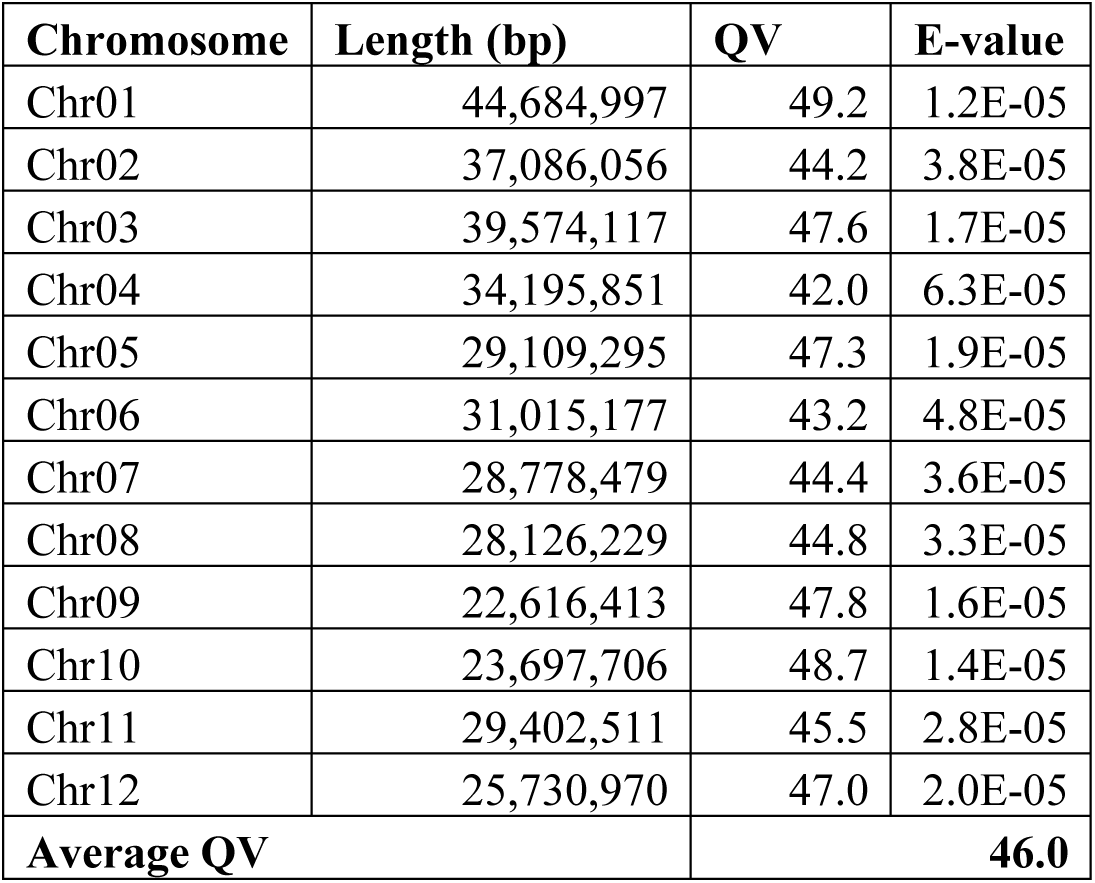
The assembly quality value (QV) of KD18 genome.

### Genome assembly validation by read mapping

We analyzed the mapping back rates of our final KD18 genome assembly using both WGS and RNA-seq data (**Table 6**). The same Illumina dataset above was mapped back to the KD18 genome assembly using Bowtie2 v2.5.5^54^ with the options *“--very-sensitive --no-unal -k 20”*, resulting an overall alignment rate of 98.07%. Additionally, a previously published low-depth resequencing dataset by Illumina from an independently sourced KD18 accession (2.9x)^16^ was also utilized to assess the released assembly. After trimming, around eight million high-quality PE reads (95.9% of original raw read pairs) were retained, of which 99.43% were mapped onto the genome. Furthermore, pooled RNA-seq data from mixed tissues of the indica rice cultivar Funong A^34^ were mapped onto the KD18 genome using HISAT2 v2.2.1^29^ with default settings, resulting in an overall mapping rate of 96.87%. Altogether, the high read mapping rates from cultivar-specific, cross-cultivar WGS and RNA-seq data suggest that our KD18 genome assembly accurately represents the sequencing data and exhibits high completeness.

**Table 6.** Mapping-back rates of WGS and RNA-seq data against the KD18 genome assembly.

|  | **KD18 deep WGS data**<br>(150×, this study) | **KD18 genome survey**<br>(2.9×, Higgins *et al.* 2021<sup>16</sup>) | **RNA-seq data**<br>(Hong *et al.*, 2026<sup>34</sup>) |
| --- | --- | --- | --- |
| **Total reads (pairs)** | 202,893,908 | 4,084,295 | 47,988,016 |
| Aligned concordantly 0 times | 5,161,864 (2.54%) | 1,462,217 (35.80%) | 2,320,163 (4.83%) |
| Aligned concordantly exactly 1 time | 128,167,491 (63.17%) | 1,700,489 (41.63%) | 44,161,512 (92.03%) |
| Aligned concordantly >1 times | 69,564,553 (34.29%) | 921,589 (22.56%) | 1,506,341 (3.14%) |
| Pairs aligned concordantly 0 times; of these: | 5,161,864 | 1,462,217 | 2,320,163 |
| Aligned discordantly 1 time | 280,844 (5.44%) | 735,981 (50.33%) | 62,234 (2.68%) |
| Pairs aligned 0 times concordantly or discordantly; of these: | 4,881,020 | 726,236 | 2,257,929 |
| Mates make up the pairs; of these: | 9,762,040 | 1,452,472 | 4,515,858 |
| Aligned 0 times | 7,833,037 (80.24%) | 46,846 (3.23%) | 3,000,589 (66.45%) |
| Aligned exactly 1 time | 694,069 (7.11%) | 261,692 (18.02%) | 1,453,176 (32.18%) |
| Aligned >1 times | 1,234,934 (12.65%) | 1,143,934 (78.76%) | 62,093 (1.37%) |
| **Overall alignment rate (%)** | **98.07** | **99.43** | **96.87** |

### Whole-genome macro-synteny with other rice cultivars

As an additional measure of genome assembly quality in regard to genome structure, we analyzed the syntenic relationships of KD18 genome with those of other rice cultivars using GENESPACE^55^ (**Fig. 5**). In this analysis, we included two indica rice genomes, MH63^8^, T197^7^, and one japonica Nipponbare genome (*Osa1_r7*, release 7, from the Rice Genome Annotation Project database - RGAP)^10^. As expected, our KD18 genome shared a high level of synteny with other rice genomes, especially with those of the indica rice. This high level of synteny with other high-quality rice genomes indicates that the KD18 genome assembly accurately preserves the expected chromosome structure and gene order.

**Figure 5.**
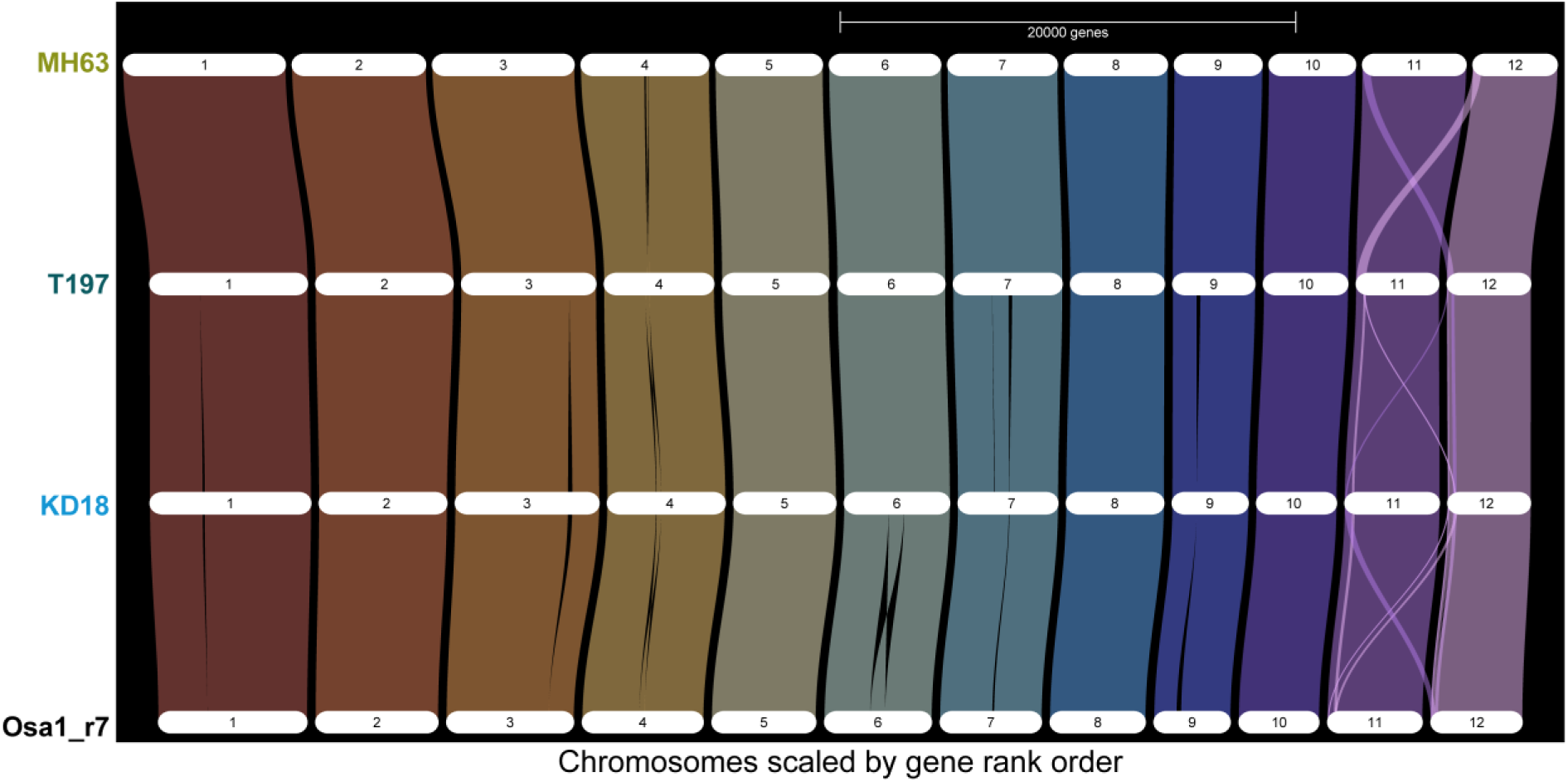
Riparian plot showing macro-syntenic regions and large-scale structural rearrangements in the genome of KD18 compared with other selected *O. sativa* genomes across 12 rice chromosomes. Chromosomes are scaled by gene rank order. The plot was generated by using GENESPACE^55^. The selected genomes include indica rice KD18 (this study), MH63^8^, T197^7^ and japonica Nipponbare (*Osa1_r7*, release 7, from the Rice Genome Annotation Project database - RGAP)^10^.

## CODE AVAILABILITY

All software packages and pipelines were executed according to the respective software documentation. Software versions and settings are provided in detail in the **Methods** section. No custom scripts or code were used in this study.

## AUTHOR CONTRIBUTIONS

T.M.V and T.Q.N conceived the project. T.M.V supervised the data generation, analysis and manuscript preparation. K.H.D.D. prepared samples, extracted DNA for both Illumina and ONT; constructed library and performed ONT sequencing. N.V.H performed genome assembly and annotation; analyzed data, wrote the first draft and revised the manuscript. T.Q.N. maintained the KD18 germplasm and performed multi-season agronomic and phenotypic characterization. All authors read, revised and approved the final version of the manuscript.

## ACKNOWLEDGEMENTS

The authors gratefully acknowledge Nguyen Tat Thanh University, Ho Chi Minh City, Vietnam, for supporting this study by providing high-performance computing resources for data analysis.

